# Disruption of the nascent polypeptide-associated complex leads to reduced polyglutamine aggregation and toxicity

**DOI:** 10.1101/2024.04.19.590245

**Authors:** Leeran B. Dublin-Ryan, Ankan K. Bhadra, Heather L. True

**Affiliations:** Department of Cell Biology and Physiology, Washington University School of Medicine, St. Louis, MO, USA

## Abstract

The nascent polypeptide-associate complex (NAC) is a heterodimeric chaperone complex that binds near the ribosome exit tunnel and is the first point of chaperone contact for newly synthesized proteins. Deletion of the NAC induces embryonic lethality in many multi-cellular organisms. Previous work has shown that the deletion of the NAC rescues cells from prion-induced cytotoxicity. This counterintuitive result led us to hypothesize that NAC disruption would improve viability in cells expressing human misfolding proteins. Here, we show that NAC disruption improves viability in cells expressing expanded polyglutamine and also leads to delayed and reduced aggregation of expanded polyglutamine and changes in polyglutamine aggregate morphology. Moreover, we show that NAC disruption leads to changes in *de novo* yeast prion induction. These results indicate that the NAC plays a critical role in aggregate organization as a potential therapeutic target in neurodegenerative disorders.

## Introduction

Protein folding is vital for cell survival and maintaining cellular homeostasis. Some proteins have amyloidogenic amino acid regions, which make them prone to misfolding and aggregation, causing disease [1]. For example, Huntington’s disease is caused by elongated repeats of glutamine (polyQ) in the Huntingtin protein that lead to misfolding and aggregation of the protein [2]. The accumulation of these Huntingtin aggregates in neurons entangles cellular resources and alters cellular processes, causing stress and eventual cell death [3–8]. Currently, there are no therapies available for Huntington’s disease, and the process by which Huntingtin protein aggregates and causes the disease is not fully understood. Many studies on the effects of the expanded polyQ region of Huntingtin protein (htt-103Q) have been conducted in yeast models, showing that polyglutamine toxicity disrupts endoplasmic reticulum (ER) stress responses, such as ER-associated degradation (ERAD) [8]. Inhibition of aggregation of mutant huntingtin (htt-103Q) is a promising strategy to slow down progress of the disease. One of the ways to do that is by modulating the activity of chaperone proteins [9].

Molecular chaperones operate as a vast and complex network, assisting in co- and post-translational protein folding. Furthermore, molecular chaperones are known to assist in ribosome shuttling to the endoplasmic reticulum and mitochondria by recognizing signal sequences on nascent polypeptides [10–12]. Many molecular chaperones are required for cellular growth and deleting these chaperones results in cytotoxicity [13–15].

Surprisingly, past research has shown that deleting components of a co-translational molecular chaperone complex, the nascent polypeptide-associated complex (NAC), rescues yeast prion-induced cytotoxicity by altering the balance and localization of other molecular chaperones [16]. The NAC is highly conserved and deletion of the complex is embryonically lethal in many multicellular organisms [13,17]. The NAC is comprised of two classes of subunits known as alpha (α) and beta (β). Egd2 is the α subunit, while in yeast there are two β subunits, Egd1 and Btt1. Btt1 arose in yeast after a genome duplication event, is known as the β’ subunit, and is 100-fold less concentrated in the cell than Egd1. The NAC is most prominently present in the cell as a heterodimer of one α subunit and one β subunit [18,19]. The β subunit reversibly binds the complex to the ribosome in a one-to-one ratio near the ribosomal exit tunnel [20]. Both α and β subunits are the first point of contact for nascent polypeptides within the exit tunnel and as they emerge from the ribosome [21]. The NAC also assists in the recognition of peptide signals and shuttling the ribosome to the ER and mitochondria [10–12]. Our group previously showed that partial deletion of the NAC rescued cytotoxicity in a yeast prion model [16]. Work from other groups has shown that overexpression of the NAC prevents aggregation of other misfolding proteins in yeast and other organisms [12,22]. Because of this work, we wanted to evaluate NAC modulation effects on human misfolding protein-induced cytotoxicity in yeast cells. Of note, loss of proteostasis is a hallmark of several of these protein-misfolding diseases. For the restoration of the proteostasis network, eukaryotic cells have developed compartment-specific stress response mechanisms. One such component is the endoplasmic reticulum (ER). The ER plays a vital role in cellular protein quality control by extracting and degrading proteins that are not correctly folded or assembled into native complexes. This process, known as ER-associated degradation (ERAD), ensures that only properly folded and assembled proteins are transported to their final destinations [23]. Stresses that disrupt ER function lead to the accumulation of unfolded proteins in the ER, a condition known as ER stress. If ER stress persists, cells activate mechanisms that result in cell death [24]. Cells adapt to ER stress by activating an integrated signal transduction pathway called the unfolded protein response (UPR) [24,25]. In the yeast *Saccharomyces cerevisiae*, this UPR pathway is mediated by Ire1 [26].

Another important aspect of co-translational protein folding is a translational control mechanism that links codon usage to protein expression levels [27]. It has been shown that the speed of ribosome movement at and around the start codon determines the rate at which subsequent ribosomes bind to the mRNA. To allow efficient rebinding of an initiating mRNA by downstream ribosomes, the actively bound ribosome must move away from the start codon quickly. Conversely, the slow movement of the bound ribosome around the start codon is sufficient to limit translation initiation rates. This occurs even when other features of the mRNA support high initiation rates. This translational control mechanism is a crucial contributing factor to expression levels in recombinant protein expression constructs, as well as determining expression levels of endogenous eukaryotic genes [27].

Herein, we report that partial deletion of the NAC improves cellular growth in yeast expressing a toxic expanded polyQ protein (htt-103Q) and reduces aggregation of polyglutamine. We also found that there was a significant increase in ubiquitination in NAC-deleted strains expressing htt-103Q. Additionally, we show that partial deletion of the NAC reduces the induction of certain yeast prion strains. These results identify the NAC as a potential therapeutic target to reduce amyloidogenic protein aggregation and further elucidate the mechanism by which NAC disruption improves the cellular environment in the face of protein misfolding stress. Curiously, we also found that deleting portions of the NAC enhances the expression of downstream UPR proteins. At the same time, these NAC deletion strains showed variability in protein expression level under a controlled codon usage variant experimental setup. These results suggest that the loss of the NAC can alter translation rates, which may improve protein folding in the cytosol (or alter aggregation propensity), while at the same time adding stress to the ER (presumably due to changes in translocation).

## Results

### NAC modulation alters the codon-dependent translation rate

Most translated nascent polypeptide chains are protected from degradation through cotranslational folding and interaction with the NAC. We have shown previously that deleting subunits of the NAC rescues yeast prion-induced cytotoxicity by altering the balance and localization of other molecular chaperones [16]. In addition, we hypothesized that protein folding was altered such that newly synthesized prion protein was slower to join aggregating conformers. It was unclear, however, whether the chaperone changes had any impact on the translation itself. Global translation (as assessed by polysome profiles) appears similar in the presence and absence of the NAC in yeast. However, there may be scenarios where co-translational folding by chaperones is required to achieve steady-state expression of a protein. Codon usage has evolved as a means to optimize translation on individual mRNAs, as well as global optimization of ribosome availability [27]. Codon decoding time is a partial predictor of protein expression levels [28]. In addition, codon optimality plays a role in protein folding, which, in turn, could alter chaperone dependence.

We wanted to test whether the absence of the NAC complex might impact protein expression through suboptimal codons. To test this, we used well-characterized controlled codon usage variants for two different protein-coding sequences. These constructs express firefly luciferase derivatives with a deletion of the last three amino acids of the native sequence which maintains full activity while removing the peroxisomal localization of the CFLuc and Renilla luciferase (RLuc) proteins [29–31]. These variants were further modified by systematically replacing codons with the slowest possible codon coding for the same amino acid to generate ‘min’ variants of the two reporter genes (minCFLuc, and minRLuc). The naturally occurring versions of these genes, which contain mixtures of fast and slow codons, were denoted as standard or ‘sta’ variants. We found that the NAC deletion strain transformed with these CFLuc and RLuc constructs showed lower luciferase activity as compared to the wild-type strain (Fig 1A), but that difference was enhanced and only significant with the ‘min’ constructs (Fig 1B). As expected, western blots showed that luciferase steady-state levels were significantly lower in NAC deletion strains as compared to wild-type (S1 Fig). These results indicate that reduced codon optimization may exacerbate the requirement for the NAC for either the expression or stability of some of the proteome.

**Fig 1.**
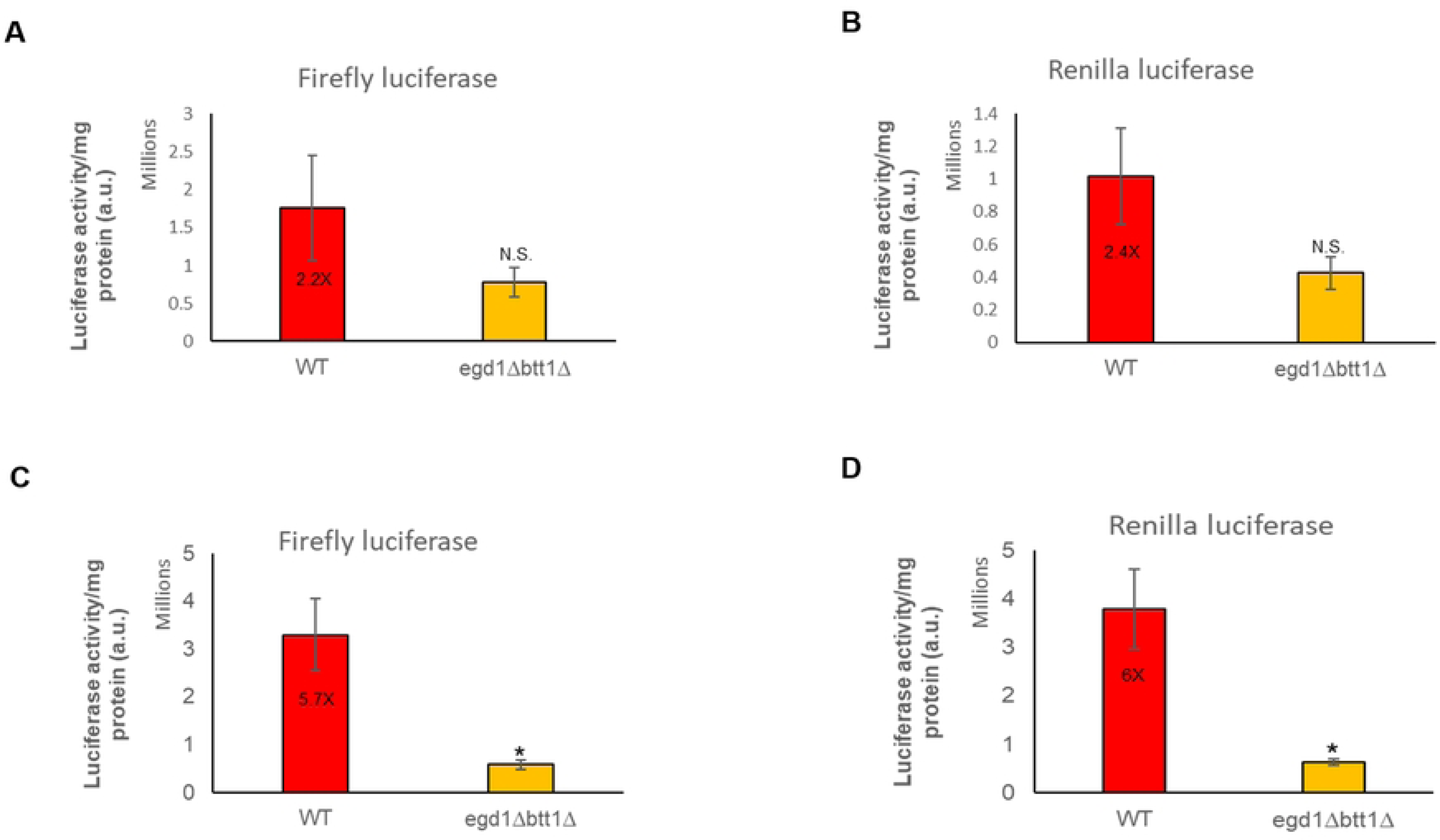
NAC modulation alters the codon-dependent translation rate. (A), (B), (C), and (D). Yeast 74-D694 WT and NAC deletion strains expressing minCFLuc, and RLuc codon variants from centromeric (single copy) plasmids using identical transcriptional and translational control sequences, consisting of the transcriptional promoter and 5′-UTR of the yeast TDH3 (glyceraldehyde-3-phosphate dehydrogenase, GPD) gene, and of the 3′-UTR and transcriptional terminator sequences of the yeast ADH1 (alcohol dehydrogenase) gene. Both TDH3 and ADH1 are highly expressed endogenous yeast genes. These are grown in SD-Ura broth. Dual luciferase assay (Promega) was performed on the same samples following the kit manufacturer protocol with minor modifications (n=3).

### NAC modulation effects on yeast expressing polyglutamine-expanded proteins

Previously, we have shown that NAC deletion rescued prion-induced cytotoxicity [16], while others have shown that NAC overexpression delayed protein aggregation [22]. Based on this work, we hypothesized that modulating NAC subunit expression would rescue toxicity induced by human misfolding proteins. To test this hypothesis, we expressed multiple disease-causing human misfolding proteins in the NAC deletion strains as well as WT yeast strains overexpressing NAC subunits. We expressed two galactose-inducible partial huntingtin proteins, htt-103Q and htt-25Q in a complete set of NAC deletion strains and a WT strain. We spotted the *egd1Δbtt1Δ, egd1Δegd2Δ, nacΔ,* and WT transformants on selective media containing glucose or galactose to evaluate toxicity induced by expanded polyglutamine (Fig 2A). We also transformed a WT 74-D694 yeast strain with plasmids expressing NAC β-subunit Egd1 or NAC α-subunit Egd2. We then transformed these strains with a plasmid expressing htt-103Q or an empty vector and spotted them to evaluate toxicity (Fig 2B). We were surprised to see the partial rescue of htt-103Q toxicity in the *egd1Δbtt1Δ* and *egd1Δegd2Δ* strains but not in strains overexpressing Egd1 or Egd2. Other work has shown that overexpression of the NAC α and β subunits suppressed expanded polyglutamine aggregation in *C. elegans* [22]. We further tested this hypothesis on multiple galactose-inducible alpha-synuclein proteins, including wild-type and two disease-causing mutants (A30P and A53T). Interestingly, we found negligible rescue in the *egd1Δegd2Δ* and *nacΔ* strains expressing alpha-synuclein WT and disease-causing mutant constructs when compared to WT expressing the same constructs (S2 Fig).

**Fig 2.**
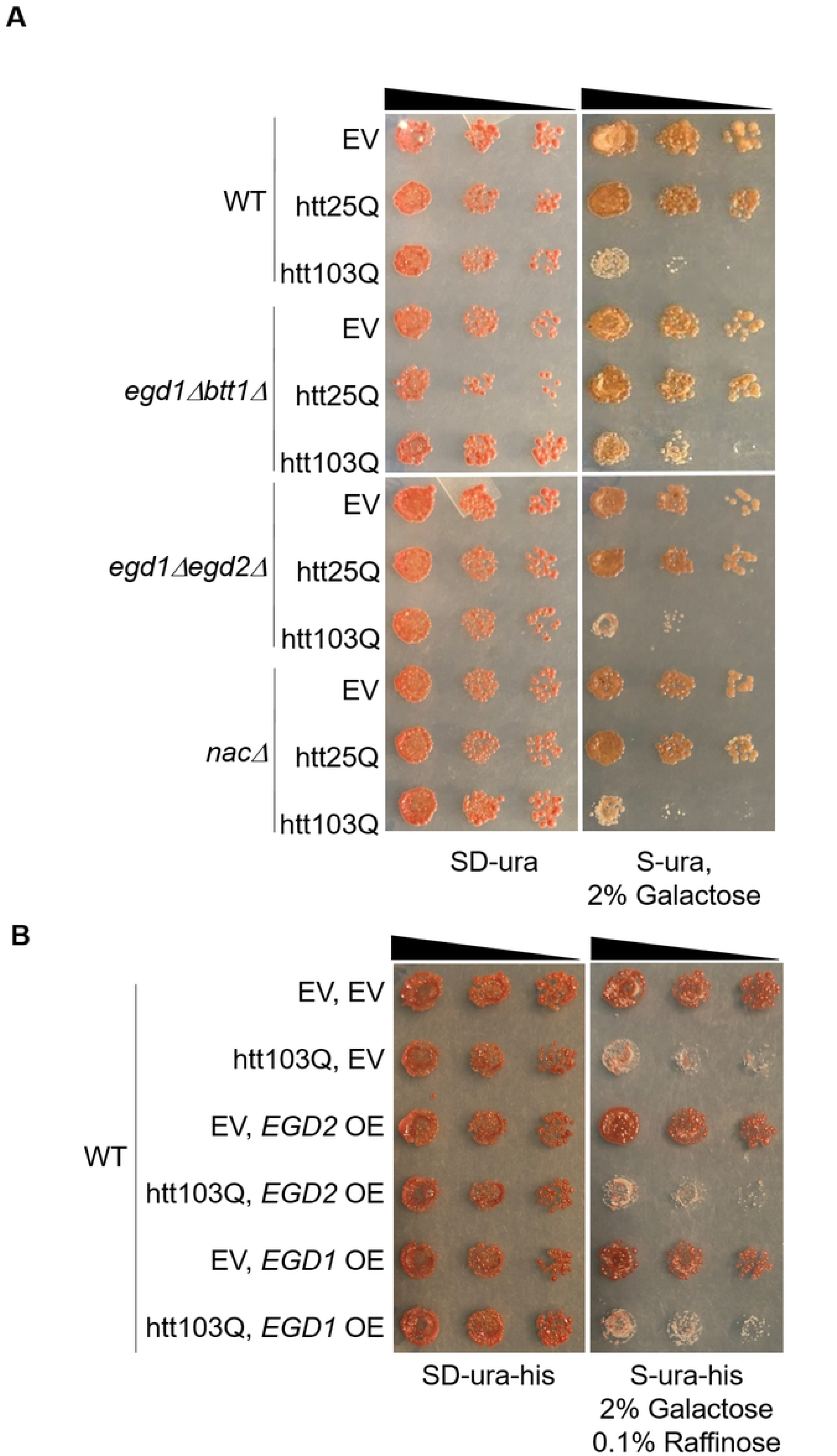
NAC modulation effects on yeast expressing polyglutamine expanded proteins. (A) Yeast 74-D694 WT and NAC deletion strains expressing Gal1-inducible EV, htt25Q, or htt103Q constructs were serially diluted 5-fold and spotted onto ¼ YPD (not shown), SD-ura, and S-ura, 2% galactose (first 3 spots shown) to monitor expanded polyglutamine toxicity and the ability of nac deletion to rescue expanded polyglutamine cytotoxicity (n=3). (B) A yeast 74-D694 WT strain expressing EV, EGD2, or EGD1 constructs in combination with Gal-inducible EV or htt103Q constructs were serially diluted 5-fold on ¼ YPD (not shown), SD-ura-his, and S-ura-his, 2% galactose, 0.1% raffinose (first three spots shown) to evaluate expanded polyglutamine toxicity and the ability of EGD1 and EGD2 overexpression to rescue expanded polyglutamine cytotoxicity (n=3).

### NAC disruption delays and reduces polyglutamine aggregation

Intrigued by these results, we investigated the aggregation of htt-103Q. While the exact mechanism by which polyQ-expanded htt causes toxicity is unknown, we know the aggregation of polyQ is a contributing factor and marker of toxicity [32]. To visualize htt-103Q aggregation in *egd1Δbtt1Δ* and *egd1Δegd2Δ* we induced expression of the protein for 6 and 20 hours using a galactose inducible promoter. We then conducted microscopy and visualized the CFP-tagged htt-103Q protein. We saw significantly reduced aggregation in the *egd1Δbtt1Δ* strain as compared to WT and *egd1Δegd2Δ* at both 6 and 20 hours of induction (Fig 3A). WT cells expressing htt-103Q for 6 hours showed that an average of 37% of cells contained aggregates while only 4.5% of the *egd1Δbtt1Δ* cells contained aggregates (Fig 3B). Even after inducing expression of htt-103Q for 20 hours, 41% of WT cells contained aggregates while 13% of the *egd1Δbtt1Δ* cells contained aggregates (Fig 3C).

**Fig 3.**
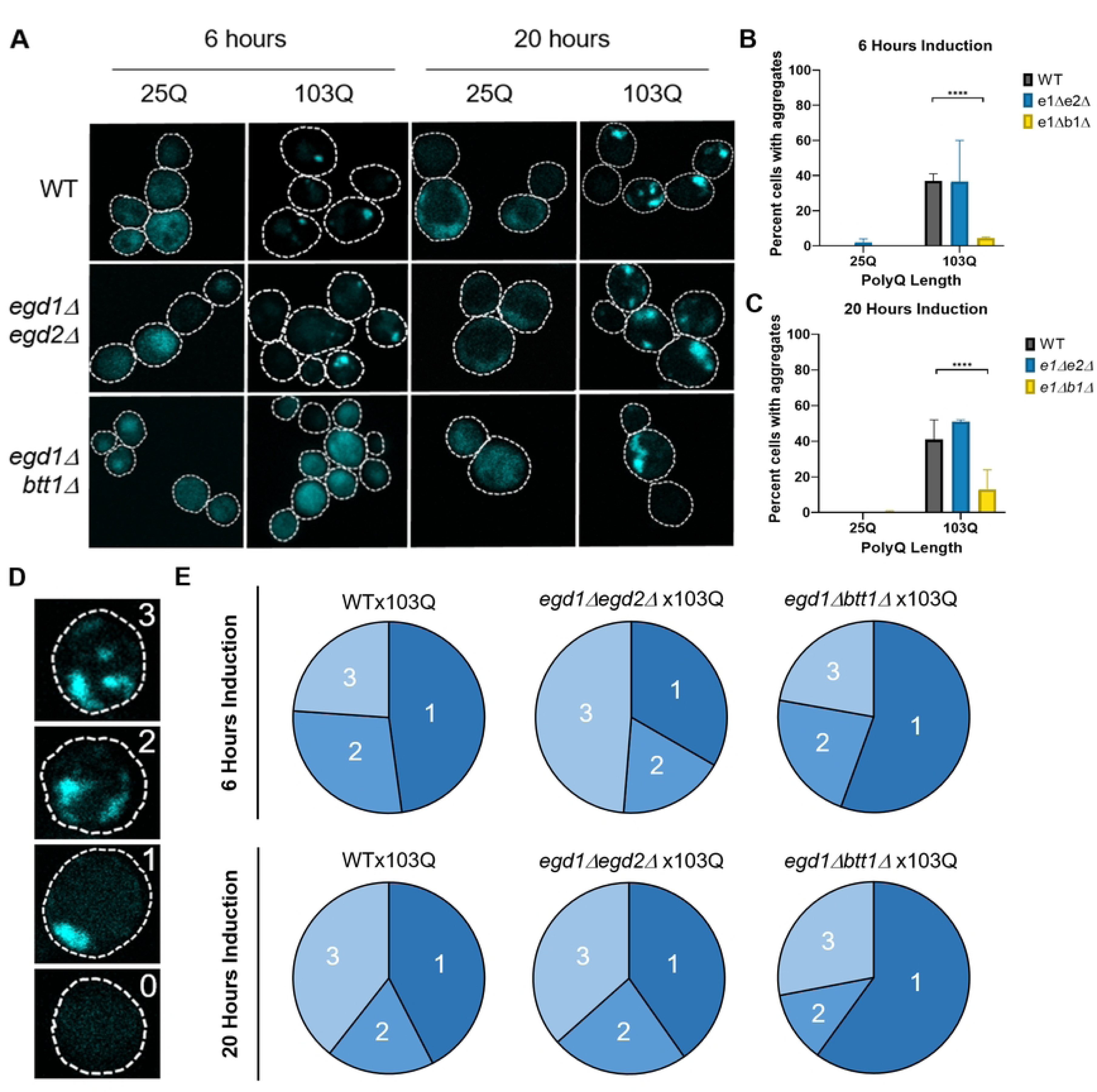
NAC disruption delays and reduces polyglutamine aggregation and changes aggregate distribution. (A) WT and NAC deletion strains expressing Gal1-htt25Q-CFP or Gal1-htt103Q-CFP constructs were grown in selective media in the presence of 2% galactose for 6 and 20 hours. Two-dimensional z-stack images were taken with a Zeiss LSM 880 Airyscan Two-Photon Confocal Microscope with a 63x oil immersion objective. The *egd1Δbtt1Δ* strain showed a smaller population of cells with aggregates than the WT or *egd1Δegd2Δ* strains at both time points. (B) Quantification of percent cells expressing Gal1-htt25Q-CFP or Gal1-htt103Q-CFP constructs containing aggregates after 6 hours induction by 2% galactose in selective media. Significance was determined by Fisher’s exact test (****=<0.0001). (C) Quantification of percent cells expressing Gal1-htt25Q-CFP or Gal1-htt103Q-CFP constructs containing aggregates after 20 hours induction by 2% galactose in selective media. Significance was determined by Fisher’s exact test (****=p value<0.0001). (D) Representative images of cells with 0, 1, 2, or 3 aggregates. Microscopy of WT and NAC deletion strains expressing Gal1-htt25Q-CFP or Gal1-htt103Q-CFP constructs for 6 and 20 hours were evaluated based on this scale. (E) WT and NAC deletion strain cells expressing Gal1-htt103Q-CFP constructs for 6 or 20 hours and containing aggregates were evaluated as having 1, 2, or 3 or more aggregates. The resulting population distributions are represented as percentages in pie graphs.

### NAC disruption changes polyglutamine aggregate distribution

Interestingly, we saw morphological differences between the htt-103Q aggregates present in the mutant strains as compared to the aggregates in wild-type cells. To quantify aggregate phenotype, we evaluated each cell expressing htt-103Q and ranked them as 0-3 depending on the number of aggregates present in the cell (Fig 3D). These analyses were of keen interest, as many groups have hypothesized and shown that soluble polyglutamine oligomers are cytotoxic, while large insoluble, polyglutamine aggregate structures may pose less risk to cellular homeostasis [7,33–35]. At 6 hours of htt-103Q induction, the aggregate populations in WT and *egd1Δbtt1Δ* aggregate populations were similar, as cells containing one aggregate comprise about half of the aggregate population in both strains (Fig 3E). However, cells containing three or more aggregates appear in 49% of cells with aggregates in the *egd1Δegd2Δ* strain at the 6-hour induction time point (Fig 3E). At 20 hours htt-103Q induction, aggregate populations observed in WT and *egd1Δegd2Δ* are similar (cells with one aggregate: 42% and 40%, two aggregates: 18% and 23%, and three or more aggregates: 39% and 37%) (Fig 3E). Conversely, the most abundant aggregate population in the *egd1Δbtt1Δ* strain after 20 hours of induction is one aggregate, which makes up 60% of the aggregate population in *egd1Δbtt1Δ* cells (Fig 3E).

### NAC disruption alters [*PSI*^+^] variant induction

We wanted to delve deeper into the change in aggregate morphology we saw with microscopy. We utilized the well-characterized [*PSI*^+^] prion model in yeast, as *de novo* [*PSI*^+^] strain induction can be used to better evaluate aggregate morphological differences as a result of chaperone deletion [36]. Furthermore, we wanted to investigate whether NAC disruption would change the aggregation morphology of another protein (Sup35) compared to WT. *Saccharomyces cerevisiae* harbors a wide array of prions, which are normally non-cytotoxic [37]. One prion, [*PSI^+^*], has been used to develop a colorimetric assay that allows for the evaluation of prion induction and prion strain determination [38].

Like polyglutamine, proteins that misfold to form prions can fold and aggregate in different conformations, creating different prion strains [39]. These strains can be differentiated by a variety of assays and, in the yeast [*PSI^+^*] prion system, by colony color [40]. Sup35 (yeast eRF3) is a protein involved in stop codon recognition with Sup45 [37]. Sup35 misfolds and aggregates to form the yeast prion [*PSI*^+^] [37]. The 74-D694 yeast strain contains the *ade1-14* allele; *ADE1* encodes a protein that is involved in the adenine biogenesis pathway, and *ade1-14* contains a premature stop codon [41]. When Sup35 is natively folded the premature stop codon in *ade1-14* is recognized and adenine biosynthesis is interrupted [41]. This results in the accumulation of a red byproduct in cells and results in their inability to grow on media lacking adenine [41]. When Sup35 misfolds and aggregates there is readthrough of the premature stop codon in *ade1-14*, resulting in completion of the adenine biogenesis pathway, no accumulation of red pigment in yeast cells, and colony growth on media lacking adenine [41]. [*PSI*^+^] prion strains include very weak, weak, medium, and strong [*PSI^+^*] and each strain can be recognized by the accumulation of pigment in yeast colonies, ranging from [*psi*^-^] cells appearing red and strong [*PSI*^+^] cells appearing very light or pale pink [40].

Given the observed changes in the htt-103Q aggregate morphology, we hypothesized that loss of the NAC would cause changes in the distribution of *de novo* [*PSI*^+^] prion strains when compared to *de novo* [*PSI*^+^] prion strains induced in WT yeast. To induce prion formation we expressed a Sup35 plasmid in WT and the NAC deletion strains (*nacΔ*, *egd1Δegd2Δ*, and *egd1Δbtt1Δ*) and grew the resulting transformants overnight in 1M KCl selective media. 1M KCl was selected because it has been shown to cause osmotic stress [42] and induce [*PSI*^+^] [43]. We then plated the transformants and determined the [*PSI^+^*] strain induced for a minimum of 100 colonies per yeast strain. The WT and *nacΔ* strains showed similar [*PSI^+^*] strain distributions (Table 1), however, both the *egd1Δegd2Δ* and *egd1Δbtt1Δ* strains had notable increases in very weak and weak [*PSI^+^*] and decreases in medium and strong [*PSI^+^*] when compared to WT (Table 1). While not perfectly correlative, this shift reflects the polyglutamine morphological differences we detected in *egd1Δbtt1Δ,* indicating partial NAC deletion leads to potentially widespread changes in protein folding.

**Table 1.** NAC disruption alters [*PSI^+^*] variant induction. WT and NAC deletion strains expressing a pEMBL-SUP35 construct were grown overnight in selective media containing 1M KCl, plated, and evaluated for [PSI+] prion strength by a previously described colorimetric assay. Over 100 colonies were evaluated from three independent experiments and ±values=SEM.

| [PSI<sup>+</sup>] strain | Wild Type | *nac*Δ | *egd1*Δ*egd2*Δ | *egd1*Δ*btt1*Δ |
| --- | --- | --- | --- | --- |
| Very Weak | 8.0±5.0% | 10.7±4.3% | 19.1±6.1% | 17.1±3.8% |
| Weak | 26.9±6.2% | 28.8±5.2% | 38.0±3.9% | 35.3±3.0% |
| Medium | 54.1±8.4% | 52.6±7.9% | 40.7±3.2% | 41.6±3.7% |
| Strong | 11.0±2.6% | 8.0±1.1% | 2.3±2.3% | 6.1±2.4% |

### NAC disruption-induced reduction of polyglutamine aggregation is not caused by reduced polyglutamine expression but correlates with increased ubiquitination

We considered the possibility that the reduction of htt-103Q aggregation in *egd1Δbtt1Δ* cells could be caused by reduced expression of htt-103Q, as reduced polyglutamine expression has been shown to reduce and reverse polyglutamine aggregate formation [44–46]. To determine htt-103Q expression we performed western blots for the FLAG-tag attached to the N-terminus of the htt-25Q and htt-103Q proteins induced in the NAC deletion strains for 6 hours. We saw no significant difference between htt-103Q expression in WT and the *egd1Δbtt1Δ* strain at 6 hours of induction (Fig 4A). We also detected htt-103Q aggregates after 6 and 20-hour induction in WT, *egd1Δbtt1Δ*, *egd1Δegd2Δ*, and *nacΔ* strains by filter trap assay. Filter trap assays allow for the visualization and semi-quantification of large protein aggregates, as these aggregates, unlike monomeric proteins, are unable to pass through cellulose acetate membranes [47]. We expected to see a reduction of aggregated htt-103Q by filter trap assay in the *egd1Δbtt1Δ* strain compared to WT at both 6 and 20 hours of induction. We saw this result at the 6-hour time point (Figs 4B and S3) but did not at the 20-hour time point (data not shown). Quantification of aggregated htt-103Q signal intensity showed a 53% percent reduction in the *egd1Δbtt1Δ* strain compared to WT after 6 hours of induction (Fig 4C).

**Fig 4.**
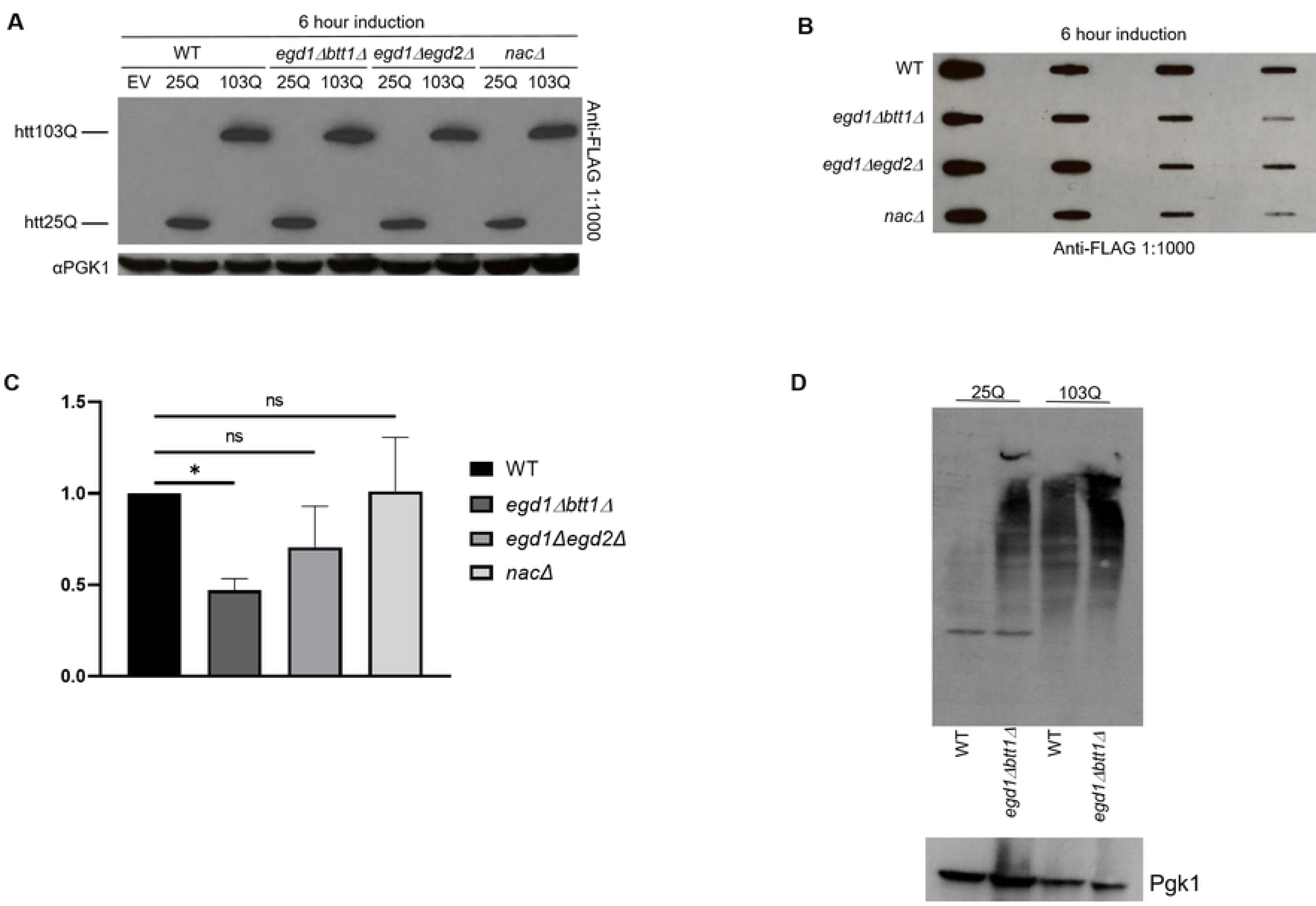
Reduction in polyglutamine aggregation in NAC-disrupted strains correlates with increased ubiquitination. (A) WT and NAC deletion strains expressing Gal1-FLAG-htt25Q-CFP or Gal1-FLAG-htt103Q-CFP constructs were grown in selective media in the presence of 2% galactose for 6 hours prior to SDS-PAGE and Western blotting for FLAG and PGK1. Western blots are representative images of three independent experiments. (B) WT and NAC deletion strains expressing Gal1-FLAG-htt25Q-CFP (not shown) or Gal1-FLAG-htt103Q-CFP constructs were grown in selective media in the presence of 2% galactose for 6 hours prior to Filter Trap Assay and Western blotting for FLAG. Western blot is a representative image of three independent experiments. (C) Quantification and normalization to WT of 3 independent Filter Trap Assay Western blots represented in 4(B), data are represented as mean ± SEM, significance was determined by paired t-test, *=p value<0.05, and ns=p value>0.05. (D) Western blot image showing the Ubiquitinated proteins in WT and *egd1Δbtt1Δ* strains co-expressing htt25Q, htt103Q constructs with an Ub-X-*LacZ* reporter: pGal-Ub-P-*LacZ*. Pgk1 showing the loading control. Representative image shown here.

One of the ways cells may counteract disease-associated expanded polyglutamine stretches is via activation of the ubiquitin-proteome system (UPS) [48–50]. It is known that polyQ expression leads to impairment of specific branches of the UPS pathway in yeast [8,49]. Also, polyQ expression in yeast triggers mitochondrial dysfunction and ER stress, which is modulated by specific UPS activities [8,51,52]. Thus, yeast polyQ models are highly useful in elucidating the role of diverse Ubiquitin-related protein degradation pathways in modulating cytotoxicity. We next tried to test the ability of our NAC deletion strains to induce the activation of the ubiquitin-proteasome pathway to mitigate the htt-103Q-mediated cytotoxicity. Indeed we found that the amount of ubiquitinated proteins was higher in NAC deletion strains expressing htt-103Q as compared to the wild-type cells expressing the polyQ protein (Fig 4D).

### Unfolded Protein Response (UPR) and ER-associated Degradation (ERAD) are altered by the loss of NAC subunits

We and others (de Alamo Plos Bio 2011) have found that loss of the NAC subunits does not result in a transcriptional response that induces the unfolded protein response (UPR). As such, we set out to determine whether there was a NAC-dependent change in the UPR in the presence of htt-103Q. Interestingly, we found that the control *egd1Δbtt1Δ* strain with an empty vector (no htt-103Q) showed increased steady-state levels of BiP (Kar2) as compared to WT. BiP, a central player in the UPR, is regulated by mechanisms other than transcription [53]. Recent studies suggest that the NAC and the signal recognition particle (SRP) bind signal peptides simultaneously [54] and that the NAC serves to provide specificity to target proteins appropriately to the ER. In the absence of the NAC, SRP promotes the mislocalization of proteins to the organelle. As such, we predicted that both the UPR and ERAD would be increased in the absence of NAC subunits. We probed cell lysates for changes in expression in SRP, ERAD, and UPR and found that the NAC deletions had increased steady-state expression of some subunits as compared to WT strains (Figs 5 and S4), indicating an abrupt activation of the unfolded protein response in these NAC deletion strains. Moreover, the toxicity of htt depends on its ubiquitination [55]. When the ER is overloaded with misfolded proteins, CDC48 extracts proteins from the ER and subsequent turnover of misfolded ER resident proteins depends on CDC48 and the ubiquitin ligase San1. This suggests that an increase in the ubiquitination of htt might be apparent in the NAC deletion strains as well. We did see an increase in overall protein ubiquitination in the deletion strains expressing both htt-25Q and htt-103Q relative to WT (Fig 4D).

**Fig 5.**
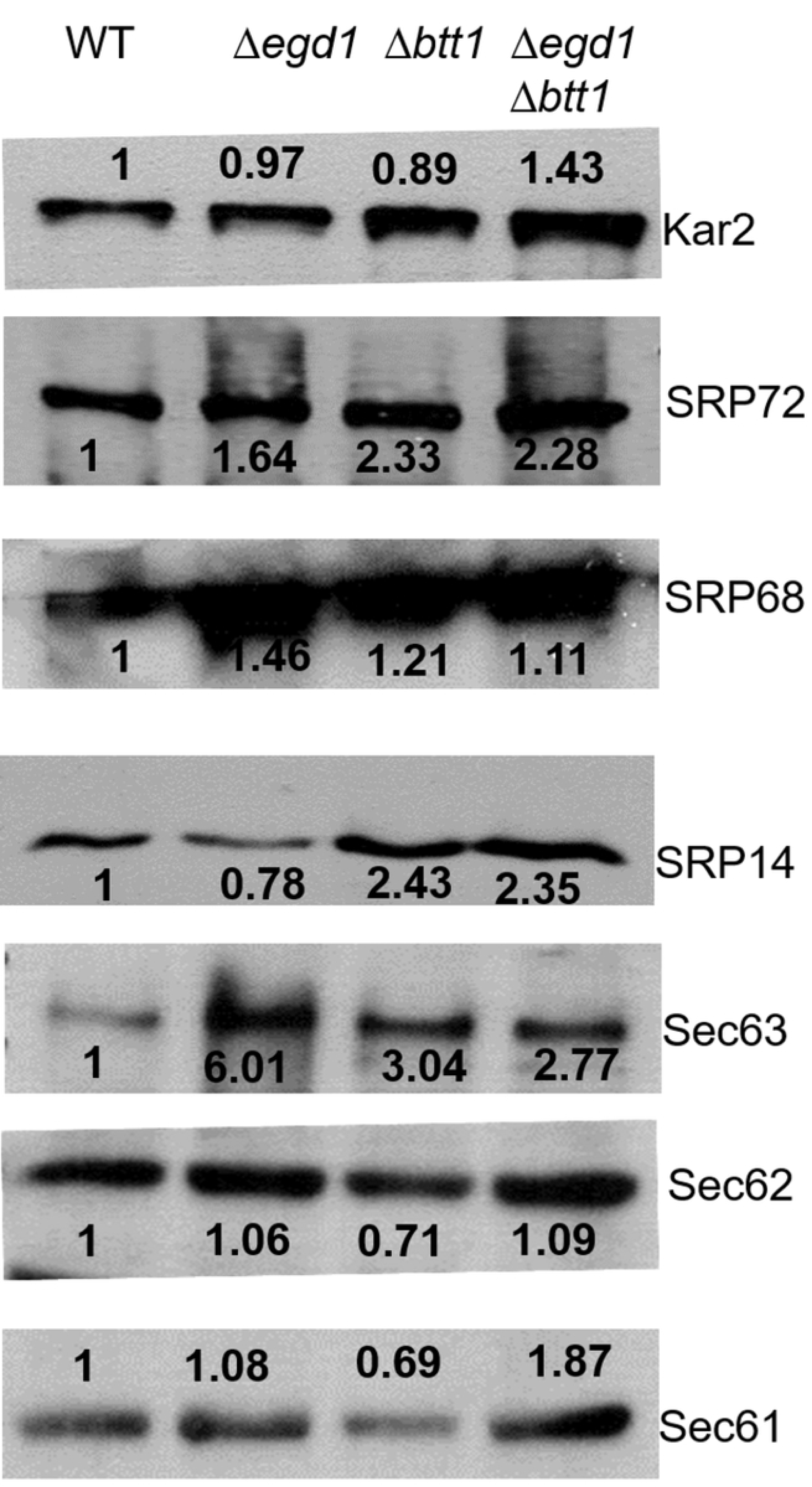
UPR and ERAD are both altered by the loss of NAC subunits. Yeast 74-D694 WT and NAC deletion strains were grown in YPD broth and lysed using standard yeast lysis protocol for western blot analysis using various Unfolded Protein Response (UPR) pathway and ERAD pathway protein antibodies. The values represent the fold difference in the expression of each protein to the WT strains, normalized to loading control (Pgk1). Representative images are shown here.

## Discussion

Our work shows that disruption of the NAC can be beneficial to cell survival in the face of protein misfolding and aggregation stress. We have provided evidence that the deletion of NAC subunits leads to improved viability in cells overexpressing htt-103Q. Furthermore, we have shown that htt-103Q aggregation is delayed in *egd1Δbtt1Δ* when compared to htt-103Q aggregation in WT, though the expression of htt-103Q is unchanged. Filter trap assays show that less htt-103Q protein is aggregated after 6 hours of induction in *egd1Δbtt1Δ* cells when compared to htt-103Q aggregated protein in WT cells. Interestingly, NAC disruption leads to changes in htt-103Q aggregate morphology as well. Lastly, we show that NAC disruption changes the distribution of [*PSI*^+^] strain inductions compared to WT. This suggests that the NAC is an important factor in aggregate formation and organization and that NAC disruption may be capable of inducing widespread alterations in aggregate morphology.

These counterintuitive results warrant further investigation. Beyond the main results showing deletion of *EGD1* and *BTT1* leads to delayed and decreased aggregation of the htt-103Q, we noticed that complete NAC deletion and deletion of *EGD1* and *EGD2* do not result in robust rescue of cellular viability when htt-103Q is expressed (Fig 2B). This indicates that individual NAC subunits may have cellular roles that are important in protein aggregation and maintaining cellular viability. Previous work showing variable rescue of canavanine toxicity suggested this as well (Keefer and True, 2016). We suggested previously that there was a delay in the joining of newly synthesized proteins to pre-existing aggregates when NAC was deleted and that this may be partly due to a change in SSB (RAC) localization and activity [16]. Here, we add to the potential mechanisms at play and suggest that the rate of translation may be altered in cells lacking the NAC. It is well known that codon bias can modulate the rate of translation [56] and that the rate of translation can impact co-translational protein folding [57–59].

Interestingly, one avenue of investigation into the rescue of htt-103Q toxicity rescue led to our discovery that some UPR and ERAD proteins are already at a higher steady-state level in the NAC deletion strains as compared to WT. This was unexpected because previous work by our group and others reported no transcriptional response using UPRE reporters. While this may not be a key feature of the toxicity rescue, it makes sense with the more recent structural studies that suggest that the NAC and SRP bind simultaneously to the signal peptide protruding from the ribosome and the absence of NAC causes more indiscriminate ER translocation [55].

Overall, this work shows that disruption of a chaperone complex has the potential as a therapeutic target in protein misfolding and aggregation disorders. Further investigation is needed to understand how the deletion of *EGD1* and *BTT1* leads to delayed and reduced htt-103Q aggregation, htt-103Q aggregate morphology changes, and altered structures in new aggregates, as seen with the change in *de novo* [*PSI*^+^] strain induction.

## Materials and methods

### Yeast strains, plasmids, cultures, and transformations

All yeast strains are derivatives of 74-D694 (*ade1-14 his3-Δ200 leu2-3,112 trp1-289 ura3-52*). Single and combinatorial genetic deletions of *EGD1, EGD2, and BTT1* were made as previously described [16]. Plasmids expressing huntingtin exon I fusions controlled by the inducible *GAL1* promoter (p416Gal1-FLAG-htt-25QΔPro-CFP and p416Gal1-FLAG-htt-103QΔPro-CFP) were kindly gifted by S. Lindquist and were made as previously described [60]. Plasmids expressing alpha-synuclein constructs controlled by the inducible *GAL1* promoter (pRS426-Gal1-SNCA^WT^-GFP, pRS426-Gal1-SNCA^A30P^-GFP, pRS426-Gal1-SNCA^A53T^-GFP) were kindly gifted by M. Jackrel. pTH726-CEN-RLuc/minCFLuc and pTH727-CEN-RLuc/staCFLuc were a gift from Tobias von der Haar (Addgene plasmid # 38210 and 38211). Egd1 and Egd2 overexpression plasmids are under the control of their endogenous promoters (p413-Egd1, p413-Egd2) and were made using standard molecular biology techniques and confirmed by sequencing.

### Antibodies

A monoclonal anti-FLAG antibody was obtained from Sigma-Aldrich. A monoclonal anti-PGK1 antibody was obtained from ThermoFisher Scientific. A rabbit anti-mouse secondary antibody was obtained from ThermoFisher Scientific. A mouse monoclonal anti-ubiquitin antibody was obtained from Santa Cruz Biotechnology. Antibodies against SRP72, SRP68, SRP14, Sec61, Sec62, Sec63, and Kar2 were produced in rabbits and are obtained as kind gifts from Prof. Peter Walter’s lab. A rabbit polyclonal anti-luciferase antibody and a goat anti-rabbit secondary antibody were obtained from Millipore-Sigma.

### Dual-Luciferase activity assay

The termination efficiency of strains transformed with the luciferase-based reporter constructs was determined in 96-well microtitre plates as described elsewhere [27] with slight modifications, using a commercial dual luciferase assay (Dual-Glo Assay, Promega, UK). Wildtype and NAC deletion strains transformed with pTH726-CEN-RLuc/minCFLuc and pTH727-CEN-RLuc/staCFLuc were inoculated into 1 ml of SC medium lacking uracil at 200 rpm at 30°C. After overnight growth, 100 µl of these cultures were transferred into 900 µl of fresh medium in a new 96-well microplate, and grown for an additional 4 h. Immediately before the luciferase measurements, 40 µl of passive lysis buffer (PLB, Promega, UK) was added per well to 150 µl culture grown from above to each well in triplicate in a 96 well plate). Culture/ PLB mixture (50 µl) was then mixed with 50 µl of the Firefly luciferase substrate in the wells of an opaque 96-well plate, incubated for 20 min at room temperature, and Firefly luciferase activity was measured in a luminometer. Stop-and-Glo reagent (50 µl) was then added per well, and the Renilla luciferase activity was measured after another 20 min of incubation.

### Prion manipulation

WT and NAC deletion yeast strains transformed with p426-Sup35 plasmid were grown at 30°C for 16 hours with rotation in 1M KCl SD-ura media to induce *de novo* prion formation. OD600 of each culture was determined and they were then plated on 150mm ¼ YPD solid media plates at dilutions that would render 200-500 colonies. These plates were then incubated at 30°C for 6 days and subsequently incubated at 4°C overnight for further color development. Each colony was evaluated for color and scored [*psi*^-^] if red and [*PSI*^+^] if pink, white, or sectored. At least 100 [*PSI*^+^] colonies were picked and pinned on ¼ YPD, SD-ade, and 5mM GdnHCl and grown for 5 days at 30°C to evaluate [*PSI*^+^] strength and color change due to [*PSI*^+^] induction. Each experiment was repeated at least 3 times.

### Fluorescent microscopy

Cells expressing p416Gal1-FLAG-htt-25QΔPro-CFP, p416Gal1-FLAG-htt-103QΔPro-CFP, or p414Gal1-EV were grown overnight in SD-ura. Cells were washed in S-ura, then grown for the indicated incubation time in S-ura, 2% galactose to induce expression of the htt-25Q or htt-103Q. Cells were imaged after 6 and 20 hours of induction using an x63 objective and cyan filter on a Zeiss LSM 880 air scan two-photon confocal microscope. Images were blinded to the researcher and aggregation of the htt-103Q was assessed phenotypically. Collected data were graphed and analyzed using GraphPad Prism version 8.1.1 for Windows, GraphPad Software, La Jolla, California USA. Results represent a compilation of data collected during 3 separate experiments.

### Yeast phenotypic and growth assays

WT and all combinatorial deletions of NAC subunits harboring p416Gal1-FLAG-htt-25QΔPro-CFP, p416Gal1-FLAG-htt-103QΔPro-CFP, pRS426-Gal1-SNCA^WT^-GFP, pRS426-Gal1-SNCA^A30P^-GFP, pRS426-Gal1-SNCA^A53T^-GFP, or p416Gal1-EV were grown overnight in SD-ura. Cultures were normalized to 0.1 OD600 and spotted in 5 5-fold dilutions onto YPD, SD-ura, S-ura, 2% galactose, 1% raffinose, S-ura, 2% galactose, 0.1% raffinose, and S-ura, 2% galactose. A WT yeast strain harboring p413-Gal1-FLAG-103Q-GFP and pRS316-Egd2 or pRS316-Egd1 was grown overnight in SD-ura-his. Cultures were normalized to 0.1 OD600 and spotted in 5 5-fold dilutions onto YPD, SD-ura-his, S-ura-his, 2% galactose, 1% raffinose, S-ura-his, 2% galactose, 0.1% raffinose, and S-ura-his, 2% galactose. Induction plates were grown for 5 days at 30°C and plates containing glucose were grown for 3 days at 30°C. Experiments were repeated at least 3 times.

### Protein analysis

Cells harboring p416Gal1-FLAG-htt-25QΔPro-CFP, p416Gal1-FLAG-htt-103QΔPro-CFP, or p416Gal1-EV were grown overnight in SD-ura, washed 2x with S-ura, and induced for 6 or 20 hours in S-ura, 2% galactose. Cell lysates were prepared as previously described. Total protein concentrations were determined by Bradford assays, normalized to ensure equal loading, and mixed with SDS-PAGE sample buffer. Samples were boiled for 10 minutes at 100°C before loading onto 10% polyacrylamide gel and run at a constant current of 100V. Blots were transferred overnight, incubated with primary and secondary antibodies, and visualized with enhanced chemiluminescence and film (GeneMate). Bands were analyzed and quantified with ImageJ and normalized to PGK1. Experiments were repeated at least 3 times.

For western blotting of downstream UPR proteins, wild-type and NAC deletion strains were grown in YPD and the cells were lysed and processed in the same manner as mentioned above. Western blotting for the codon bias experiment using an anti-luciferase antibody was performed with Wild-type and NAC-disrupted cells expressing pTH726-CEN-RLuc/minCFLuc and pTH727-CEN-RLuc/staCFLuc in SD-ura media and processed similarly as described above.

The *in vivo* UPS functionality was measured using cells of wild-type and NAC deletion strains expressing htt-25Q and htt-103Q; co-transformed with an Ub-X-*LacZ* reporter: pGal-Ub-P-*LacZ*. Post induction, cells were lysed, and a western blot was performed in the same manner as described above with an anti-ubiquitin antibody.

### Filter trap assay

The modified protocol from Sin et al., 2018 was followed. Cells harboring p416Gal1-FLAG-htt-25QΔPro-CFP, p416Gal1-FLAG-htt-103QΔPro-CFP, or p414Gal1-EV were grown overnight in SD-ura, washed 2x with S-ura, and induced for 6 hours in S-ura, 2% galactose. Cell lysates were prepared as previously described. Lysates were normalized to 500ug total protein in Filter Trap Assay Buffer and serially diluted three times at 1:2 dilutions. The resulting 2μ filter was incubated with primary and secondary antibodies, and visualized with enhanced chemiluminescence and film (GeneMate). Bands were analyzed and quantified with ImageJ. Ponceau staining was used as a loading control. Experiments were repeated at least 3 times.

## Acknowledgements

We thank Drs: M. Jackrel, Peter Walter, Tobias von der Haar, and S. Lindquist for the reagents. We thank Rachel Bouttenout for assistance with the transposon screen and strain creation. We are appreciative of Dr. C. Weihl, and M. Pullen for their helpful discussions and comments on this work.

## Supporting information

**S1 Fig. NAC deletion strains showed lower luciferase steady-state levels** Yeast 74-D694 WT and NAC deletion strains expressing minCFLuc, and RLuc codon variants from centromeric (single copy) plasmids using identical transcriptional and translational control sequences, consisting of the transcriptional promoter and 5′-UTR of the yeast TDH3 (glyceraldehyde-3-phosphate dehydrogenase, GPD) gene, and of the 3′-UTR and transcriptional terminator sequences of the yeast ADH1 (alcohol dehydrogenase) gene. Both TDH3 and ADH1 are highly expressed endogenous yeast genes. These are grown in SD-Ura broth and lysed using standard yeast lysis protocol for western blot analysis using Luciferase antibody.

**S2 Fig. NAC modulation effects on yeast expressing *α*-synuclein proteins.** Yeast 74-D694 WT and nac deletion strains expressing Gal1-inducible EV or WT, A30P, or A53T α-synuclein constructs were serially diluted 5-fold and spotted onto ¼ YPD (not shown), SD-ura, and S-ura, 2% galactose (first 3 spots shown) to monitor α-synuclein toxicity and the ability of nac deletion to rescue α-synuclein cytotoxicity (n=3).

**S3 Fig. Ponceau staining shows similar protein concentrations loaded in filter trap assay.** WT and NAC deletion strains expressing Gal1-FLAG-htt25Q-CFP (not shown) or Gal1-FLAG-htt103Q-CFP constructs were grown in selective media in the presence of 2% galactose for 6 hours prior to Filter Trap Assay and Western blotting for FLAG. After developing 2μ filter was soaked in Ponceau staining to show total protein load. All strains are expressing Gal1-FLAG-htt103Q-CFP.

**S4 Fig. Pgk1 blots showing loading controls used for calculations of fold change between WT and NAC deletion strains.** Representative images shown here.

**S1 Table. Plasmids used in this study.** The left most column indicates the True Lab log number of each plasmid, the second from the left column contains the plasmid names, the third from the left column indicates the auxotrophic marker of each plasmid, and the right most column identifies the control plasmids.

**S2 Table. Strains used in this study.** The left most column indicates the True Lab log number of each strain and the right column contains relevant strain information.

